# Quantification of long-term doxorubicin response dynamics in breast cancer cell lines to direct treatment schedules

**DOI:** 10.1101/2021.05.24.445407

**Authors:** Grant Howard, Tyler A. Jost, Thomas E. Yankeelov, Amy Brock

## Abstract

While acquired chemoresistance is recognized as a key challenge to treating many types of cancer, the dynamics with which drug sensitivity changes after exposure are poorly characterized. Most chemotherapeutic regimens call for repeated dosing at regular intervals, and if drug sensitivity changes on a similar time scale then the treatment interval could be optimized to improve treatment performance. Theoretical work suggests that such optimal schedules exist, but experimental confirmation has been obstructed by the difficulty of deconvolving the simultaneous processes of death, adaptation, and regrowth taking place in cancer cell populations. Here we present work characterizing dynamic changes in sensitivity to the chemotherapeutic doxorubicin in three breast cancer cell lines subjected to treatment schedules varying in concentration, interval between pulse treatments, and number of sequential pulse treatments. Cell populations are monitored longitudinally through automated imaging for 600-800 hours, and this data is used to calibrate a family of cancer growth models derived from the bi-exponential model which characterizes resistant and sensitive subpopulations. We identify a model incorporating both a period of growth arrest in surviving cells and a delay in the death of chemosensitive cells which outperforms the original bi-exponential growth model in Akaike Information Criterion based model selection, and use the calibrated model to quantify the performance of each drug schedule. We find that the inter-treatment interval is a key variable in determining the performance of sequential dosing schedules and identify an optimal retreatment time for each cell line which extends regrowth time by 40%-106%, demonstrating that the time scale of changes in chemosensitivity following doxorubicin exposure allows optimization of drug scheduling by varying this inter-treatment interval.

## Introduction

Cancer is the second most prevalent cause of death in the United States, and acquired chemoresistance is a common cause of treatment failure in cancer(1,2). While many studies have investigated the biochemical mechanisms of chemoresistance, predicting the onset of resistance and the population dynamics of sensitive and resistant cells remains a challenge(3–6).

Early chemotherapy treatment schedules, based on the approach of maximally-tolerated dose to achieve maximal killing of tumor cells, assumed a homogenous, exponentially growing cell population(7,8). Subsequent studies have pointed to the significance of intratumor heterogeneity in cell growth rate and drug sensitivity(9–18). Norton and others established that tumor kill is proportional to growth rate(19–21), and adjuvant chemotherapy schedules for breast cancer were revised to decrease the interval of therapy for fast-growing TNBC(22–24). Other studies point to metronomic therapy and adaptive therapy as potential improvements for breast cancer scheduling(25–29).

Mathematical modeling of drug responses has developed optimal solutions for drug dosing under a variety of model assumptions(30–33), highlighting the opportunity to improve cancer treatment by optimizing drug schedules. However, these efforts have generally been purely theoretical. Under the necessarily simplified assumptions of these drug sensitivity models, it is possible to find a true mathematical optimum, but these studies have not explored the magnitude or form of deviation between their models and experimental observations. This reveals a need for methods to experimentally test the response to drug schedules, and to use those experimental results to inform and calibrate models of drug response in cancer. Robust predictive models of drug response are a necessary step towards the ultimate goal of patient-specific predictive models of therapy-response and relapse.

A challenge in modeling chemoresistance is that the state of drug sensitivity is often represented as binary – sensitive or resistant. Clinically, resistance is usually inferred at very coarse time intervals – a patient is retrospectively assessed as sensitive or resistant to a course of treatment as a whole. In *in vitro* work, cell lines are labeled as sensitive or resistant as well, based on their stable drug sensitivity(34,35). This approach may obscure key underlying characteristics of the system if drug sensitivity is heterogeneous in the cancer cell population. In addition, cells may change drug sensitivity over time or in response to environmental conditions, adding temporal heterogeneity(36–41). The consequence is that cells display a distribution of responses to drug exposure and this distribution may vary with time(42–50). Experimental work describing the distribution of drug sensitivity in cancer cell populations, and the temporal changes arising from cell plasticity, is key to understanding drug resistance at the population level.

In this work, we seek to quantify the dynamic changes in drug sensitivity following treatment to iteratively optimize treatment schedules in a series of *in vitro* experiments subjecting three breast cancer cell lines to a series of pulsed drug perturbations which vary in drug concentration, inter-treatment interval, and number of serial drug exposures. Long-term automated time-lapse microscopy enables quantitation of population dynamics in multiple replicates, doses and regimens over days to weeks. Here we measured up to 12 individual culture wells in 3 cell types treated with a range of 9 doses and 13 regimens.

By calibrating the resulting data to a mathematical model, we quantify the distribution of the underlying populations with respect to model parameters of interest including resistant fraction, relapse growth rate, and sensitive cell death rate. Additionally, we characterize the dynamics of both the relapse and dying populations. In this study, we sought to capture the fact that sensitive cells do not respond to drug treatment by immediately undergoing cell death and resistant cells do not respond by continuing to proliferate at the same rate as untreated cells. Rather, the processes of cell death, cell cycle arrest, and cell growth after treatment are complex and these time scales may overlap—with some cells continuing to die in response to drug treatment, while others are simultaneously recovering from a period of arrest and beginning to proliferate.

Model selection is used to identify three phenomenological models, from a family of 18 related models which vary in the dynamics defining cell growth and death in treated populations, which most accurately parameterize the data. Time-lapse microscopy data is used to calibrate the three phenomenological models, and select the model which best characterizes the data. The optimal model is then used to quantify the impact of the tested drug schedules on cancer cell population, and identify schedules with superior performance. This demonstrates a method for optimizing drug schedules using mathematical models of drug sensitivity which are experimentally calibrated using high throughput in-vitro experiments. This method could be extended to additional drugs, drug combinations, and cancer types.

### Model description

We used a simple mathematical model that treats cancer cell population dynamics as a process consisting of a decreasing drug-sensitive subpopulation of tumor cells, and an independently growing drug-resistant subpopulation. This model has successfully described tumor dynamics in non-small cell lung cancer, melanoma(51), multiple myeloma(52), and ovarian cancer(53). After exposure to drug, the net fitness of the drug-sensitive (*S*) cells is represented in terms of an exponential decay or “tumor cell kill” rate *k*, and the net fitness of the resistant (*R*) cancer cells as an exponential growth rate *g_r_* > 0, and the initial fraction of drug-resistant cells by 0 ≤ *fr* ≤ 1. Additionally, the resistant cancer cells may experience a period of proliferation arrest *t_r_* ≥ 0. (The *R* and *S* designations are solely with respect to the specific drug exposure being examined, and are not intended to specify any particular mechanism of action.) To analyze the response of a population of *S* and *R* cells to drug treatment, we use a system of two ordinary differential equations (1 and 2) coupled by the total cell number (equation 3) to describe the dynamics of the *S* and *R* subpopulations.

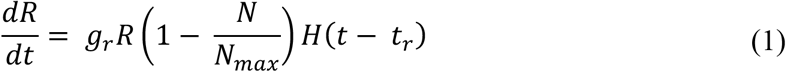

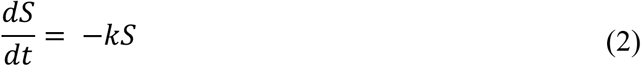

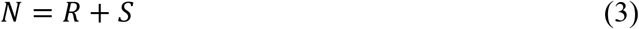

Initial conditions are specified using the resistant fraction, *f_r_*, in equations 4 and 5. *H*(*x*) is the Heaviside step function, which is 0 for negative values of the argument and 1 for positive values of the argument.

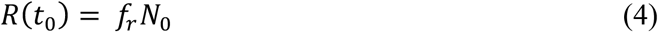

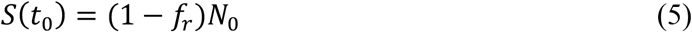

To better understand how growth and death dynamics impact drug sensitivity of the overall cell population, we use automated image analysis of time lapse microscopy data to calibrate three families of phenomenological models of the population-level drug response. Resistant cells experience a period of growth arrest, *t_r_*, followed by logistic growth at rate, *g_r_*. Sensitive cells die at rate *k*, which is subject to variation in the doxorubicin response(54), and three models for *k* are considered: 1) a time-delayed model of *k*, such that the value varies between cell growth at the initial growth rate of *g_0_* and cell death at a maximum rate of *k_d_* with exponential decay from *g_0_* to *k_d_* at a time constant of *t_d_* (equation 6), 2) a time-delayed model of *k* such that the value varies linearly between cell growth at the initial growth rate of *g_0_* and cell death at a maximum rate of *k_d_* over a total time of *t_d_* (equation 7), 3) a constant value of *k* (equation 8). Models 1 and 2 each include 6 total parameters, while the simpler model 3 includes 5.

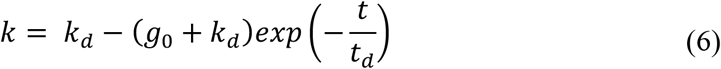

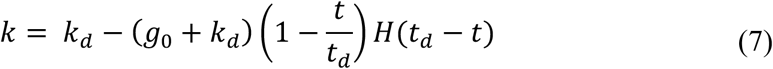

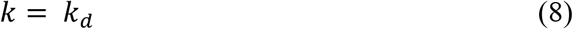

**Table 1.** Summary of model parameters

| Symbol | Model Parameter Description | Parameter Assignment |
| --- | --- | --- |
| $f_r$ | Resistant fraction | Calibrated |
| $g_r$ | Relapse growth rate | Calibrated |
| $t_r$ | Proliferation delay | Measured via clustering analysis |
| $k_d$ | Sensitive cell death rate | Calibrated |
| $t_d$ | Time constant for death delay | Calibrated |
| $g_0$ | Pre-treatment growth rate | Calibrated |
| $N_{max}$ | Carrying capacity | Calibrated |

Model 1 represents a case in which cells stochastically transition from proliferating to dying. At the time of treatment, the cell population is still proliferating (represented in the model as a negative net death rate for this cell compartment). The net death rate smoothly transitions from proliferation to death and approaches the maximum value of *k_d_* exponentially at a time constant of *t_d_*. Model 2 represents a case where cells take time, *t_d_*, to halt proliferation, again transitioning from initially proliferating at a rate of *g_0_* at the time drug exposure begins, to an eventual maximum death rate of *k_d_*. Model 3 represents the simplest case with the minimum number of parameters; here the population responds to drug exposure rapidly enough on the time scale of the experiment for transition time to be negligible. These are nested models, and we will evaluate the need for the additional parameters, which increase the complexity of the models, using model selection criteria when fitting the data to the models.

**Fig 1.**
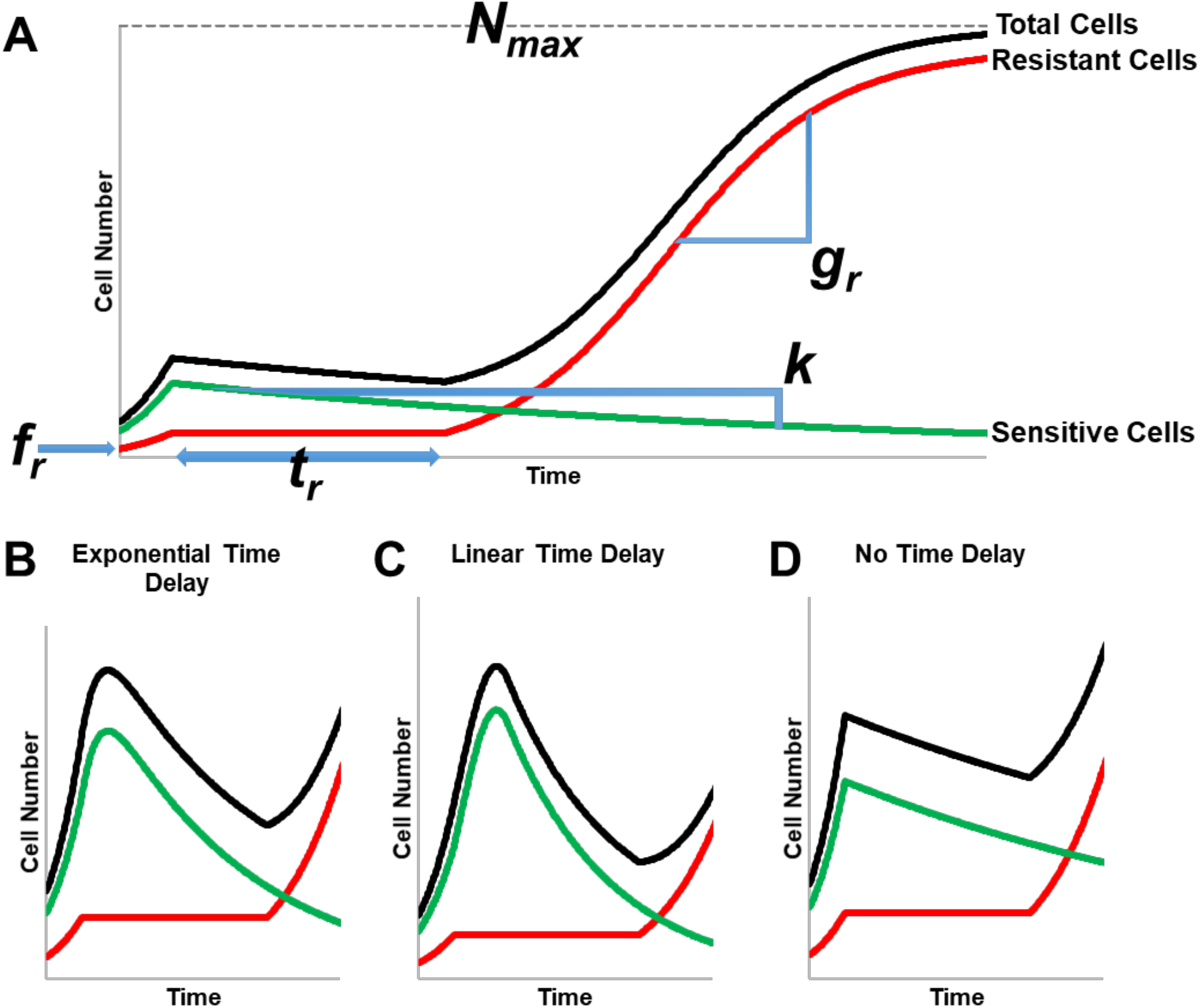
Model structures. illustrated with curves for total cell number (black) partitioned into resistant (red) and sensitive (green) fractions. Regions of the curves are marked to show the effects of key model parameters (A). The effects of three different models of *k* are shown for model 1 with an exponential time delay on *k* (B), model 2 with a linear time delay on *k* (C) or model 3 with no time delay on *k* (D).

### Model identifiability

To test the ability of our modeling framework to accurately extract parameter values and select among models, each model within the family described above was used to calibrate simulated data. This model family includes three sets of assumptions concerning the form of *k*, which are described as models 1, 2, and 3; independently, *t_r_* can be zero or non-zero. Six simulated data sets, labeled A through F and each containing 1000 time series of cell number, were generated from these six sets of assumptions concerning the underlying ground truth. For each data set, parameter values were randomly generated from within a physiologically reasonable parameter space (details provided in **Supporting Information, model identifiability results**), a cell number vector was generated based on those parameter values, and Gaussian random noise was added to each data point of the cell number vector to simulate measurement error. We were interested in determining whether the underlying model could be identified from the cell number data alone, or whether the *t_r_* value must also be specified to obtain accurate parameter values. Additionally, we sought to determine whether it was possible to reduce the number of free parameters by fixing *N_max_*. This resulted in a total of 18 models: *k* selected from models 1, 2, or 3, *t_r_* calibrated, assumed to be 0, or given as input, and *N_max_* either calibrated or fixed. Each data set was used to calibrate the 18 models, summarized in **Table 2**.

**Table 2.** Summary of model assumptions. Pearson’s correlation coefficient (PCC) *f_r_* for each model calibrated with data generated based on a matching ground truth for *t_r_* and the form of *k*.

| Model | Assignment of $t_r$,<br>proliferation delay in $R$<br>cells | $k$, death delay in $S$<br>cells | $N_{max}$ | PCC |
| --- | --- | --- | --- | --- |
| 1 | known and given as input | Exponential | Calibrated | 0.99 |
| 2 | known and given as input | Linear | Calibrated | 0.99 |
| 3 | known and given as input | None | Calibrated | 0.99 |
| 4 | calibrated | Exponential | Calibrated | 0.34 |
| 5 | calibrated | Linear | Calibrated | 0.28 |
| 6 | calibrated | None | Calibrated | 0.25 |
| 7 | 0 | Exponential | Calibrated | 0.93 |
| 8 | 0 | Linear | Calibrated | 0.94 |
| 9 | 0 | None | Calibrated | 0.99 |
| 10 | known and given as input | Exponential | Fixed | 0.74 |
| 11 | known and given as input | Linear | Fixed | 0.72 |
| 12 | known and given as input | None | Fixed | 0.71 |
| 13 | calibrated | Exponential | Fixed | 0.27 |
| 14 | calibrated | Linear | Fixed | 0.27 |
| 15 | calibrated | None | Fixed | 0.24 |
| 16 | 0 | Exponential | Fixed | 0.23 |
| 17 | 0 | Linear | Fixed | 0.29 |
| 18 | 0 | None | Fixed | 0.68 |

To check the ability of each model to extract accurate parameter values from the simulated data sets, Pearson’s correlation coefficients (PCC) between the known and calibrated values for the model parameters were computed for each combination of model and simulated data set (**Table S3**). Comparing PCC values for each model 1 through 9 to its matching model 10 through 18 reveals that calibration of the carrying capacity is necessary. Use of a fixed value for carrying capacity significantly reduces the ability of the modeling framework to accurately identify parameter values (see **Fig S3**). Consequently, models 10 through 18 were excluded from further analysis

The model sets 1, 4 and 7; 2, 5, and 8; and 3, 6, and 9 each vary solely in the handling of *t_r_*, the period of growth arrest after drug exposure. Models 4, 5, and 6 extracted parameter values with a PCC of 0.34 or less even when matched to data sets generated from matching model assumptions, while models 1, 2, and 3 each extract parameter values with a PCC of 0.99. Closer consideration revealed that the *t_r_* and *f_r_* values are not uniquely identifiable in data with any amount of noise (see **Fig S2**). Based on this difference in accuracy, models 4, 5, and 6 were excluded from further analysis. Models 7, 8, and 9, in which *t_r_* = 0, extract parameter values with PCC 0.93 or greater from data sets D-F, generated from matching model assumptions, but with PCC of 0.20 or less when calibrated to data sets A-C generated with *t_r_* ≠ 0 (**Table S3**). The key question, which must be resolved experimentally, is whether *t_r_* is in fact non-zero, as computational work alone proved unable to determine this in the analysis stage. Examination of the raw data demonstrated that the period of proliferation arrest, *t_r_*, can be non-zero, and in some cases is substantial (**Fig 2**). Consequently, models 7-9 were removed from consideration, with models 1-3 included for future analysis.

**Fig 2.**
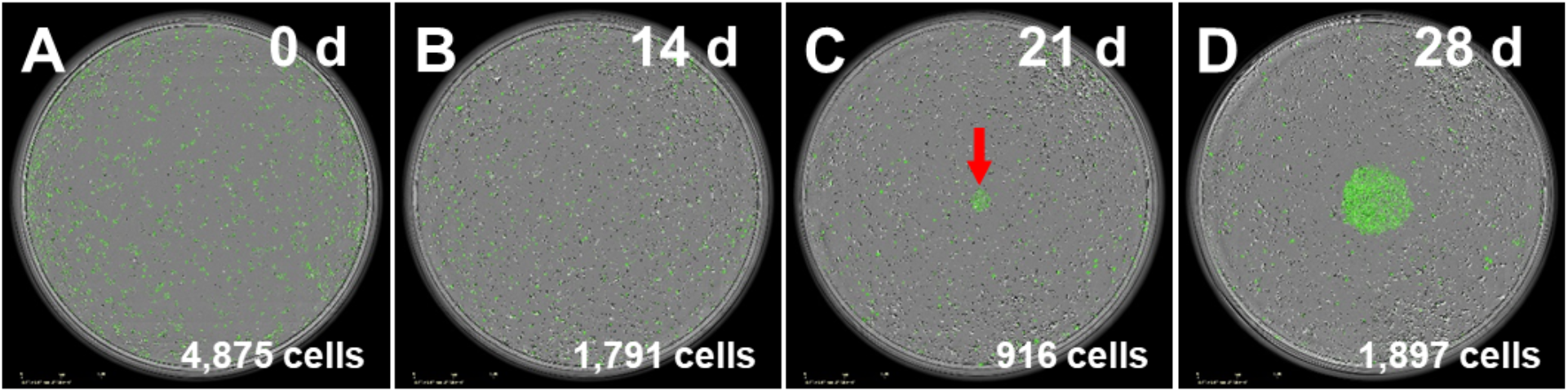
Micrograph example illustrating doxorubicin induced proliferation delay. In this example demonstrating a doxorubicin induced proliferation delay, after exposure to 150 nM doxorubicin at time 0 (A) MCF7 cells stop proliferating and remain in arrest for 315 hours. At 14 days (B), cells have died off in the intervening time, and proliferation has not occurred, and at 21 days (C), proliferation has produced a small patch of cells indicated by the red arrow; proliferation then continues and produces a patch of hundreds of cells by 28 days (D). Untreated cells grow to confluence in approximately four days.

The Akaike information criterion (AIC) was applied to the calibrations generated by models 1-3 on simulated data sets A-F to test its ability to identify the underlying model structure in cases where the ground truth is known independently. When the AIC is computed for three otherwise equivalent models which vary in the form of *k*, it selects the model which matches the ground truth of the simulated data in 97% of cases where there is no death delay, in 87% of cases if the death delay has an exponential form, and in 84% of cases when the death delay has a linear form. In each case, the correct model is identified in a majority of simulated data sets, demonstrating the utility of the AIC at identifying the model which best matches the ground truth.

## Results

### Quantification of long term population level response

To determine the dynamics of the long-term cell population response to doxorubicin, breast cancer cell populations were subjected to one to five sequential doxorubicin exposures, with the drug concentration, interval between drug exposures, or number of drug exposures allowed to vary. Time lapse microscopy and automated image analysis was used to quantify the cell number throughout the treatment period and up to four weeks after the final dose.

The long term response to a 24 hour doxorubicin pulse varies not only with drug concentration (**Fig 3A, 3D, 3G**), but also with the cell population’s history of previous exposure, even if the total dose delivered remains constant (**Fig 3B-C, 3E-F, 3H-I**). Population behavior varies over time, and the response observed at any single time point (such as 48 or 72 hours, which are frequently chosen as endpoints for single time-point observations) does not accurately characterize the overall dynamic response. Visual assessment of these cell number curves offers some intuition on the relative strength of the drug treatments: A cell population that is sensitive to a drug treatment may manifest in a curve that dips to a deeper minimum, or takes longer to resume net population growth, or shows slower population growth over the long term. As expected, for a single administration of doxorubicin, these effects of treatment increase with dose (**Fig 3A, 3D, 3G**). When multiple doses of the same concentration are administered, the total amount of drug increases with the number of exposures; this generates the similarly intuitive result that the impact of the treatment increases when going from one treatment to two sequential treatments (**Fig 3C, 3F, 3I**).

When the total drug exposure for each treatment group is held constant while the interval between the two exposures is varied, the cell population response curves vary considerably (**Fig 3B, 3E, 3H**). This indicates that the drug sensitivity of all three cell lines varies dynamically, such that the timing of a second drug exposure has a profound effect on the effectiveness of the second treatment. While these observations can be made on a qualitative level from the cell number curves, calibrating these experimental results to the models discussed above allows for quantitative investigation of the variation in response.

**Fig 3.**
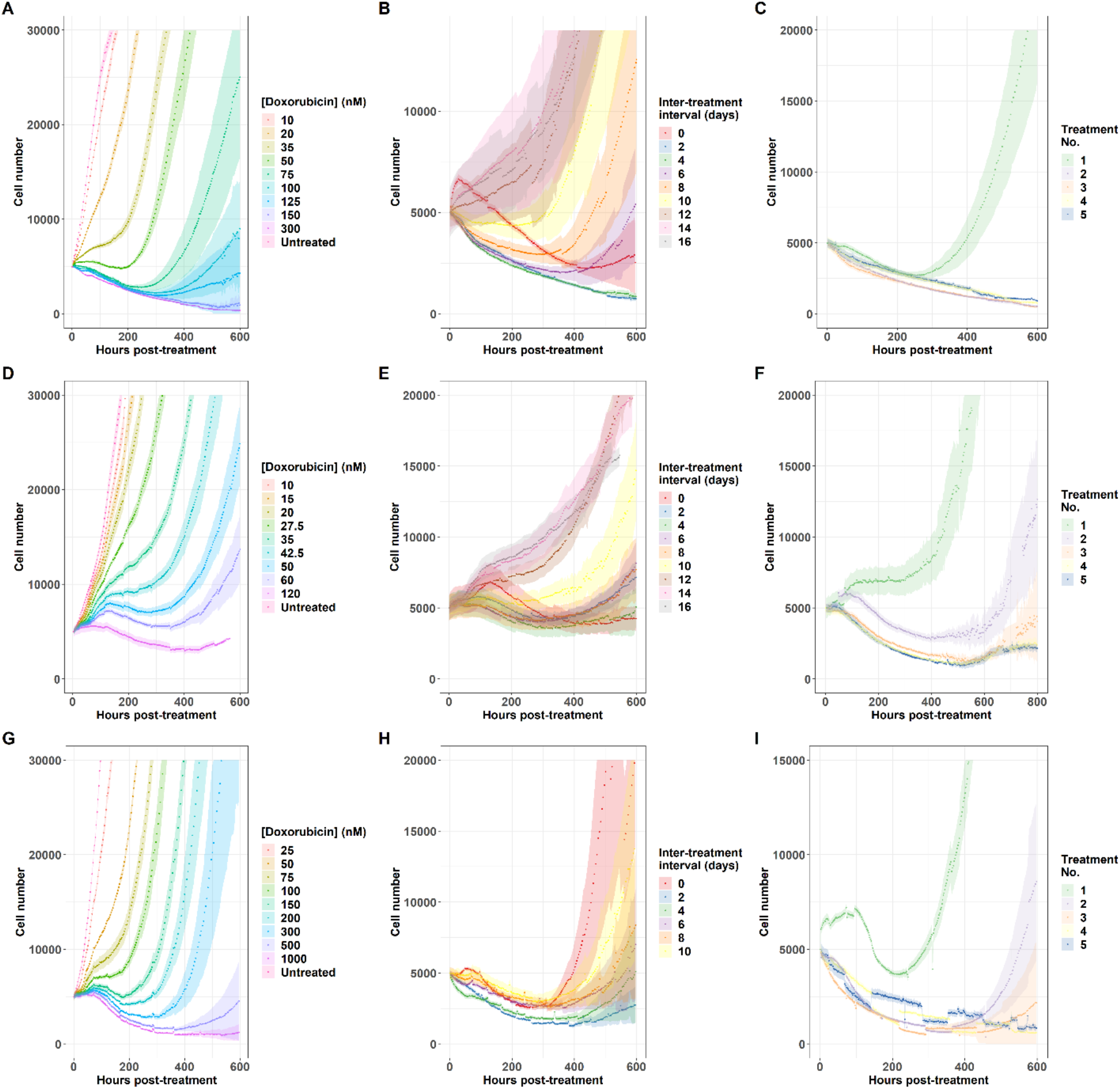
Cell population response to doxorubicin regimens. Population level response to 24 hour doxorubicin exposures is quantified as doxorubicin concentration varies in the MCF7 cell line (A), the BT474 cell line (D), and the MDA-MB-231 cell line (G), as the interval between two 24 hour doxorubicin exposures varies at 75 nM in the MCF7 cell line (B), at 35 nM in the BT474 cell line (E), and at 200 nM in the MDA-MB-231 cell line (H), and as the number of sequential 24 hour doxorubicin exposures varies at a two day interval and 75 nM in the MCF7 cell line (C), at a zero day interval (continuous exposure) and 35 nM in the BT474 cell line (F), and at a two day interval and 200 nM in the MDA-MB-231 cell line (I). Each curve represents the average of six (A, D, G) or 12 (B, C, E, F, H, I) replicate samples, and 95% confidence intervals are marked by the shaded regions. All curves are aligned such that *t* = 0 is the beginning of the final drug exposure for that treatment group.

### Fraction of non-recovering replicates

Under some treatment regimens, we observe a bifurcation in the response: some replicate cultures recover proliferative capacity during the experiment, while others do not (and therefore exhibit only cell death throughout the experiment). This necessitates split handling of the analysis as well, since the resistant population properties cannot be obtained in cases where no cells were resistant to the treatment. The fraction of replicate cultures which do not recover in each treatment group is shown in Fig 4, and these wells are excluded from the analysis of resistant cell populations.

**Fig 4.**
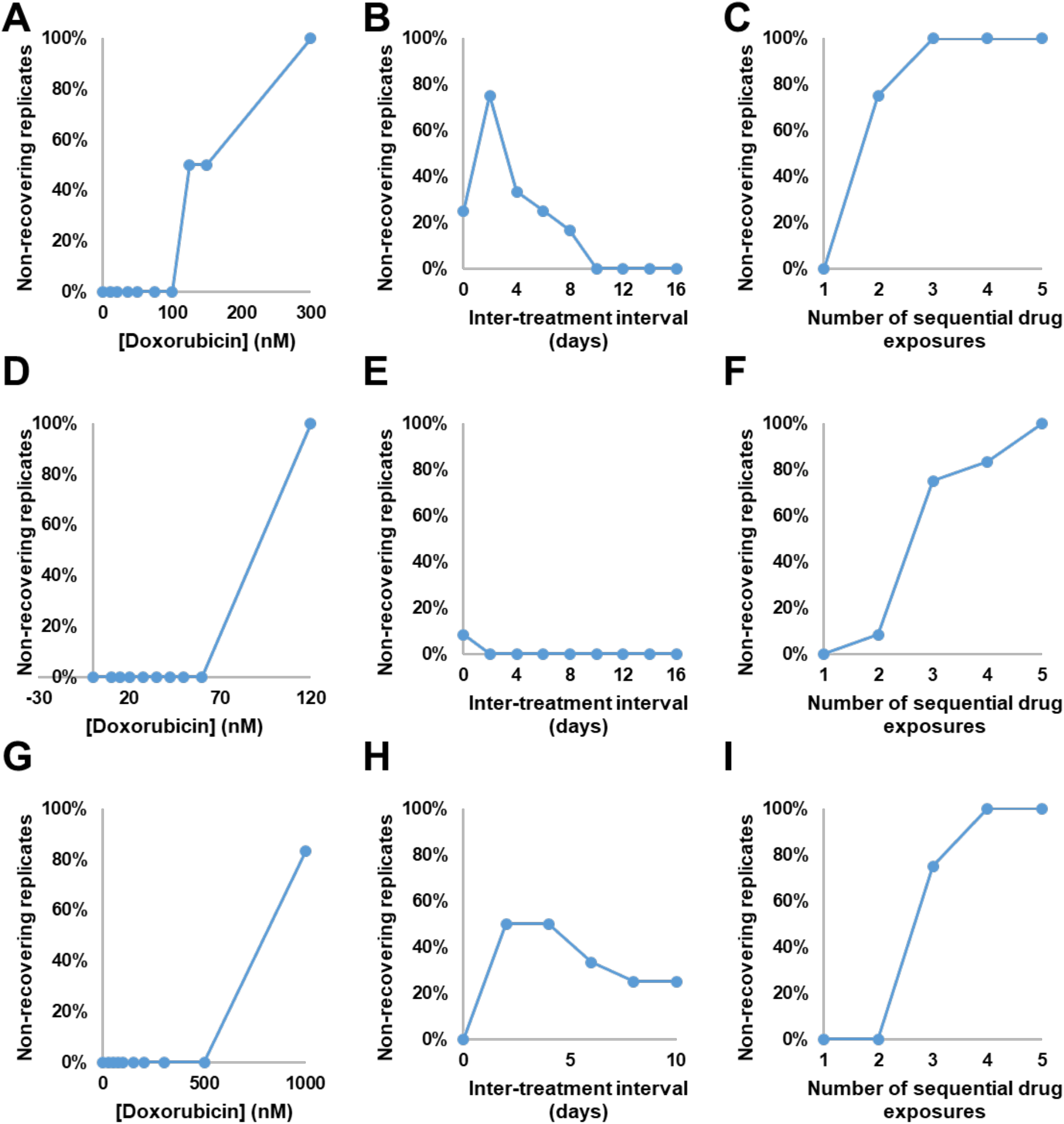
Fraction of non-recovering replicate cell populations. The percentage of replicate cultures which did not recover during the course of the experiment is shown as doxorubicin concentration varies in the MCF7 cell line (A), the BT474 cell line (D), and the MDA-MB-231 cell line (G), as the interval between two 24 hour doxorubicin exposures varies at 75 nM in the MCF7 cell line (B), at 35 nM in the BT474 cell line (E), and at 200 nM in the MDA-MB-231 cell line (H), and as the number of sequential 24 hour doxorubicin exposures varies at a two day interval and 75 nM in the MCF7 cell line (C), at a zero day interval (continuous exposure) and 35 nM in the BT474 cell line (F), and at a two day interval and 200 nM in the MDA-MB-231 cell line (I). Each value is the percentage of 6 (A, D, G) or 12 (B, C, E, F, H, I) replicates.

Although this phenomenon hinders analysis of some resistant cell properties, it also reveals which treatment regimens are capable of eliminating the entire experimental cancer cell population. The bifurcation indicates that these treatments should not be assumed to be completely effective against the cancer cell population; rather, it is sufficient for resistant cells to be rare enough in the population as a whole that the initial population of approximately 2,000 seeded cells is too low to ensure that a resistant cell will be present. Despite this caveat, these conditions do represent the practical limits of treatment regimens that can be investigated in the 96 well format. In the MCF7 cell line, doxorubicin concentrations of 125 nM or higher may result in non-recovering replicates (**Fig 4A**), and two or more sequential treatments at 75 nM may result in non-recovering replicates (**Fig 4C**). In the BT474 cell line, a doxorubicin concentration of 120 nM resulted in no recovering wells (**Fig 4D**), and two or more sequential exposures to 35 nM doxorubicin resulted in some non-recovering wells (**Fig 4F**). Similarly, in the MDA-MB-231 cell line, doxorubicin at a concentration of 1 μM (**Fig 4G**) or three or more exposures to 200 nM doxorubicin (**Fig 4I**) may result in non-recovering replicates. While the concentration thresholds vary from cell line to cell line, in each of these cases the fraction of non-recovering wells increases with the total drug exposure within a cell line.

When the inter-treatment interval is varied with the total drug exposure held constant, we find that the fraction of non-recovering wells peaks in the MCF7 cell line at 75% (nine replicates out of 12) with a two day interval (**Fig 4B**), then declines to 0% at intervals of 10 or more days. In the BT474 cell line, 8% (one replicate out of 12) fails to recover in the zero interval (continuous treatment) group, with all wells recovering at longer intervals (**Fig 4E**). In the MDA-MB-231 cell line, the fraction of wells which do not recover peaks at 50% (six out of 12 replicate wells) at an interval of 2 or 4 days, and declines at longer intervals (**Fig 4H**). In each cell line this again suggests that the drug sensitivity of these populations varies dynamically over this period of approximately two weeks after an initial drug exposure.

### Proliferation delay, *t_r_*

The model identifiability analysis demonstrates that, if a period of growth arrest is present in experimental data, then it must be incorporated into the growth model to obtain accurate parameter values *via* model calibration (**Fig S2**). Growth arrest does occur under some of the treatment regimens investigated in this series of experiments. Density based clustering analysis with manual correction for common errors is used to identify the restart of proliferation in replicate cultures which experience a period of growth arrest (**Fig 5**). When the concentration of doxorubicin in a single exposure is varied, a threshold is observed with concentrations of 50 nM or higher inducing growth arrest in the MCF7 cell line (**Fig 5A**), concentrations of 42.5 nM of higher inducing growth arrest in the BT474 cell line (**Fig 5D**), and concentrations of 150 nM or higher inducing growth arrest in the MDA-MB-231 cell line (**Fig 5G**). In the MCF7 cell line, when the interval between two 75 nM doxorubicin exposures is varied, the period of growth arrest decreases as the inter-treatment interval increases (**Fig 5B**), with intervals of 0-4 days experiencing over 200 hours of growth arrest and with some replicates at intervals of 8 or more days experiencing no growth arrest at all; since an initial 75 nM exposure results in approximately 125 hours of growth arrest, this suggests dynamic variation in drug sensitivity over this period of approximately two weeks. Likewise, in the BT474 cell line, as the interval between two 35 nM doxorubicin exposures is varied the period of growth arrest decreases with intervals of 10 days or longer resulting in no growth arrest (**Fig 5E**). Similarly, when the MDA-MB-231 cell line is exposed to two 200 nM drug pulses, the period of growth arrest is elevated for at least 10 days, with a peak of over 400 hours with a two day interval between exposures (**Fig 5H**). When the number of sequential drug exposures at a two day interval is varied, the period of growth arrest increases significantly after a second treatment in the MCF7 cell line (**Fig 5C**), the BT474 cell line (**Fig 5F**), and the MDA-MB-231 cell line (**Fig 5I**).

**Fig 5.**
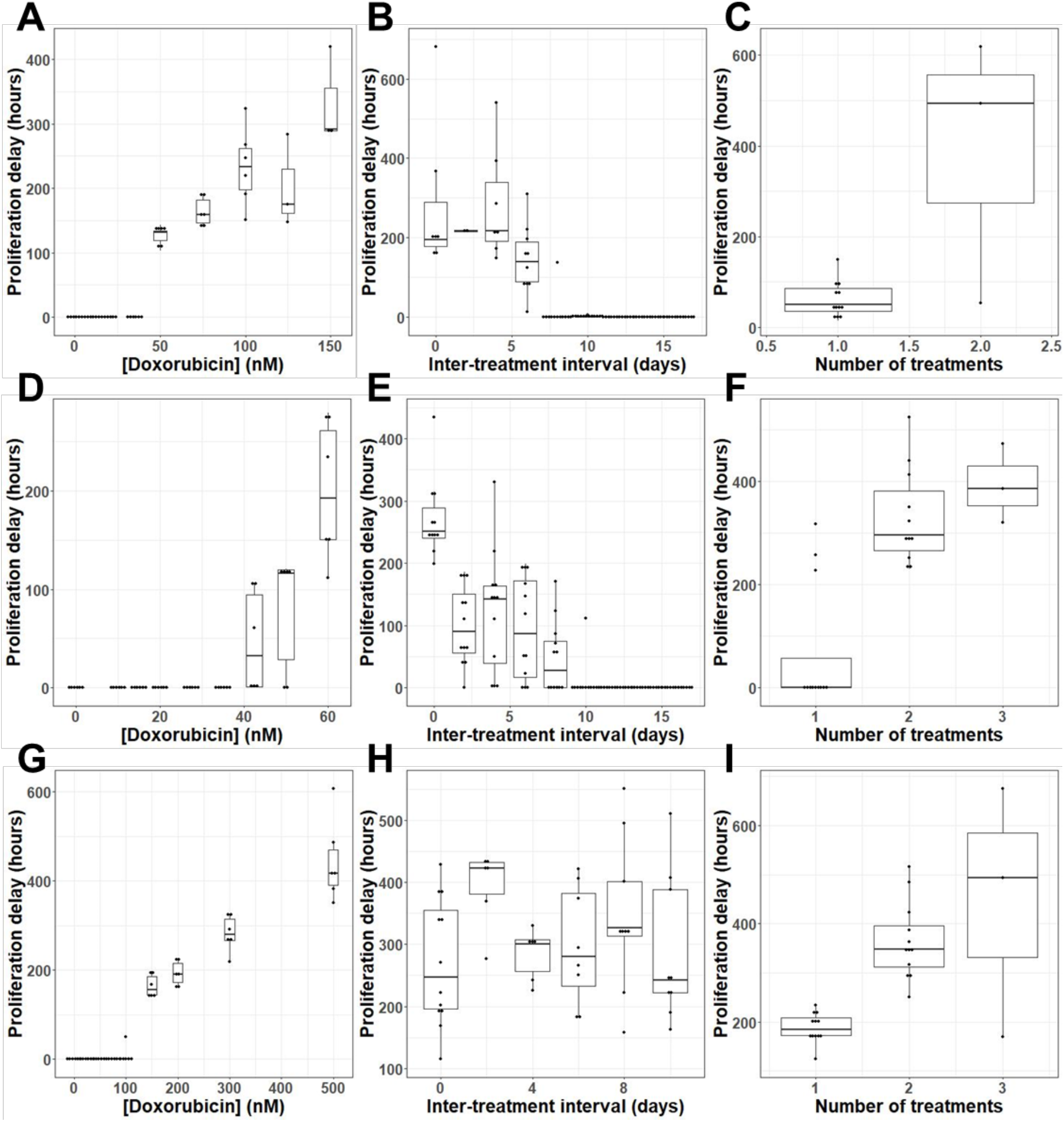
Proliferation delay, *t_r_* varies with dosing regimen. Length of growth arrest of resistant cells (*t_r_*) is displayed as a Tukey box-and-whiskers plot as doxorubicin concentration varies in the MCF7 cell line (A), the BT474 cell line (D), and the MDA-MB-231 cell line (G), as the interval between two 24 hour doxorubicin exposures varies at 75 nM in the MCF7 cell line (B), at 35 nM in the BT474 cell line (E), and at 200 nM in the MDA-MB-231 cell line (H), and as the number of sequential 24 hour doxorubicin exposures varies at a two day interval and 75 nM in the MCF7 cell line (C), at a zero day interval (continuous exposure) and 35 nM in the BT474 cell line (F), and at a two day interval and 200 nM in the MDA-MB-231 cell line (I). The number of replicates in each group varies between 3 and 12; all replicates in which recovery is observed are included (see Fig 4).

### Model selection

AIC values are calculated for each replicate culture for models 1, 2, and 3; the AIC allows comparison of the goodness of fit for models with varying numbers of free parameters. Across 648 replicate cultures calibrated to models 1, 2, and 3, the AIC indicates that model 1 is optimal in 382 replicates (59.0%), model 2 is optimal in 129 replicates (19.9%), and model 3 is optimal in 137 replicates (21.1%). This suggests that model 1 achieves the best overall performance across the range of conditions analyzed; however, models 2 and 3 are selected in a substantial minority of cases, and a more detailed investigation of the structure of these model preferences is available in the Supporting Information, structure of model preferences section (see **Fig S5** and **Fig S6**). Critical time projections and model validation results are constructed using model 1.

**Fig 6.**
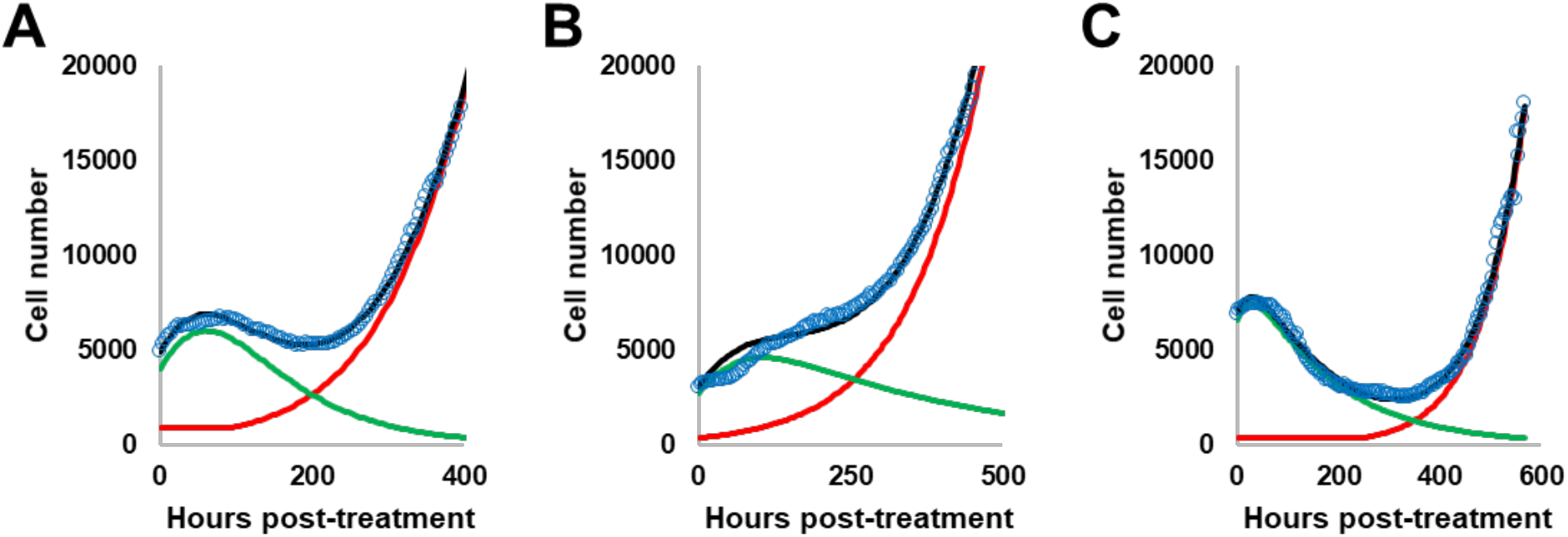
Examples of model fitting demonstrate the ability of model 1 to capture long term dynamics. Examples of model fitting are shown for a single replicate culture in the MCF7 (A), BT474 (B), and MDA-MB-231 (C) cell lines. The curve represents a response to the doxorubicin concentration used in serial treatment within the cell line – 75 nM for the MCF7 line, 35 nM for the BT474 cell line, and 200 nM for the MDA-MB-231 cell line. Data is shown as open blue circles, while the best-fit curve for model 1 is shown in black, with the resistant and sensitive compartments shown in red and green respectively.

### Model validation

A leave-one-out cross validation is used to test the predictive capability of model 1. Across 143,093 total points evaluated, 80.6% fall within the 95% confidence interval. These results are summarized in **Table 3**, with a more detailed breakdown in **Table S4** in the supporting information.

**Table 3.** Leave-one-out cross validation. The performance of model 1 at predicting data excluded from the training set is broken down by cell line and whether the replicate culture recovered or not (see Figure 4).

| Condition | Points<br>Evaluated | Within<br>95% CI |
| --- | --- | --- |
| Overall | 143093 | 80.6% |
| All cell lines, recovering wells | 94824 | 89.8% |
| All cell lines, dying wells | 48269 | 62.5% |
| MCF7 cell line, total | 42442 | 88.0% |
| MCF7 cell line, recovering wells | 25992 | 94.5% |
| MCF7 cell line, dying wells | 16450 | 77.7% |
| BT474 cell line, total | 56923 | 75.1% |
| BT474 cell line, recovering wells | 41391 | 87.3% |
| BT474 cell line, dying wells | 15532 | 42.4% |
| MDA-MB-231 cell line, total | 43728 | 80.7% |
| MDA-MB-231 cell line, recovering wells | 27441 | 89.1% |
| MDA-MB-231 cell line, dying wells | 16287 | 66.4% |

Model performance varies between cell lines and with recovery status. Model 1 performs best in the MCF7 cell line (88% of data in the 95% confidence interval), less well in the MDA-MB-231 cell line (80.7% of data in the 95% confidence interval), and worst in the BT474 cell line (75.1% of data in the 95% confidence interval). Wells which recover are better modeled (89.8% of data within the 95% confidence interval) than wells which do not (62.5% of data within the 95% confidence interval). These results indicate that model 1 captures a significant portion (but not all) of the variation in the population level response to doxorubicin.

### Critical time projection

After doxorubicin treatment, the cancer cell populations varied in rate of cell number decrease, lag time to net re-growth, and re-growth rate (Fig 3). Some cell populations remain quiescent and never initiate re-growth within the time of the experiment (Fig 4). To allow more precise and consistent comparison across the range of conditions explored in these experiments, we define a cell population as having recovered from the drug perturbation at the time, referred to as the critical time, when the population reaches twice the initial cell number for that replicate culture. This definition prioritizes the long term regrowth of a culture over its immediate response; for example, a culture which experiences a small decline in total population, but very slow regrowth, could be identified as having a longer recovery time than a culture which experiences a steep drop in population but very rapid regrowth. Utilizing a definition of cancer cell recovery or progression based on the initial population has been found to avoid counterintuitive results(55). It also enables comparison of cultures which do not experience a drop in total population, but which do vary in growth rate, such as those exposed to a low concentration of doxorubicin (Fig 3A). In some cultures, regrowth to twice the original cell number is not observed before the end of the experiment. To enable the inclusion of these results, we use model projections of the critical time based on the best-fit curve of the optimal model identified by model selection.

As the concentration of doxorubicin in a single exposure increases, the critical time increases in all three cell lines (**Fig 7A, 7D, 7G**). When the interval between two 75 nM doxorubicin exposures is varied in the MCF7 cell line (**Fig 7B**), and when the interval between two 200 nM doxorubicin exposures is varied in the MDA-MB-231 cell line (**Fig 7H**), a peak in the critical time is observed for both cell lines at a two day inter-treatment interval; this suggests that the cell population is sensitized to retreatment at this time. This result is consistent with the observation that 9/12 replicate cultures treated with this regimen in the MCF7 cell line and 6/12 replicate cultures treated with this regimen in the MDA-MB-231 cell line never recover (**Fig 4B, 4H**), and with the observation that the period of growth arrest is elevated at this time (**Fig 5B, 5H**). When the interval between two 35 nM doxorubicin exposures is varied in the BT474 cell line, the critical time is elevated at intervals of zero to eight days (**Fig 7E**), which is consistent with the elevated period of growth arrest observed in these replicates (**Fig 5E**). When the number of sequential treatments with 75 nM doxorubicin and a two day inter-treatment interval in the MCF7 cell line is varied, the critical time increases on a second treatment (**Fig 7C**); the critical time for a third treatment cannot be obtained because no replicates treated with this regimen recovered (**Fig 4C**). When the number of sequential treatments with 35 nM doxorubicin and no inter-treatment interval is varied in the BT474 cell line, the critical time increases for the second and third treatment (**Fig 7F**), and cannot be obtained for the fourth and fifth treatment because too few replicate wells recovered. Similarly, when the number of sequential treatments with 200 nM doxorubicin and a two day inter-treatment interval is varied in the MDA-MB-231 cell line, the critical time increases after a second and third treatment, and cannot be obtained for the fourth treatment because no replicates treated with that regimen recovered (**Fig 5I**).

**Fig 7.**
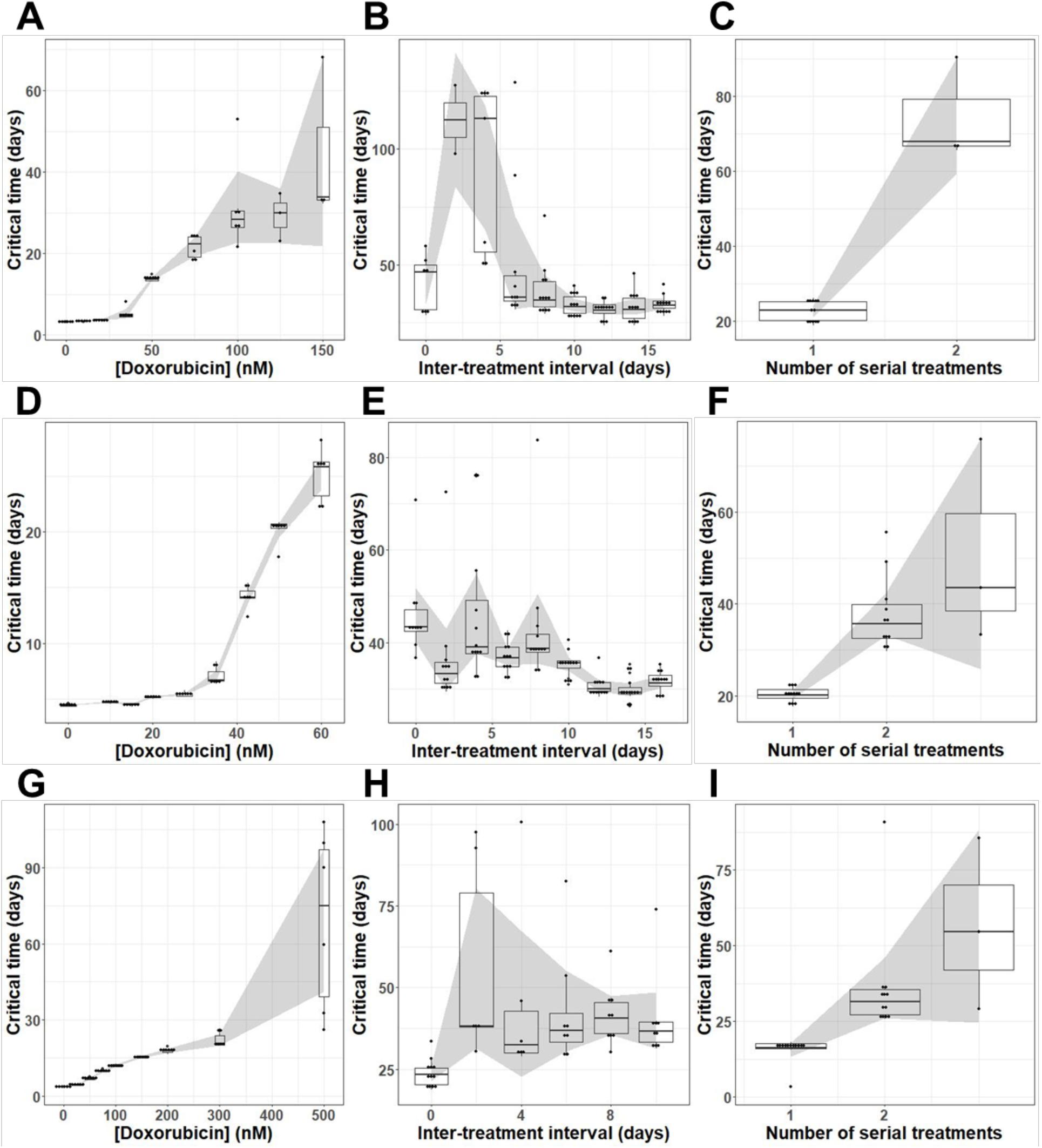
Critical time projections. Critical time is estimated based on the best fit to model 1 for each replicate culture in the MCF7 cell lines and shown as a Tukey box-and-whiskers plot as doxorubicin concentration varies in the MCF7 cell line (A), the BT474 cell line (D), and the MDA-MB-231 cell line (G), as the interval between two 24 hour doxorubicin exposures varies at 75 nM in the MCF7 cell line (B), at 35 nM in the BT474 cell line (E), and at 200 nM in the MDA-MB-231 cell line (H), and as the number of sequential 24 hour doxorubicin exposures varies at a two day interval and 75 nM in the MCF7 cell line (C), at a zero day interval (continuous exposure) and 35 nM in the BT474 cell line (F), and at a two day interval and 200 nM in the MDA-MB-231 cell line (I). The number of replicates in each group varies between 3 and 12; all replicates in which recovery is observed are included. The 95% confidence interval on the mean is shown by the shaded region.

## Methods

### Cell Culture

MCF7 (ATCC HTB-22) cells were cultured in Minimum Essential Media (MEM) (Gibco), supplemented with 10% fetal bovine serum (FBS) and 1% penicillin-streptomycin (P/S) (Gibco); these cells were maintained at 37° C in a 5% CO_2_ atmosphere(56).

BT474 (ATCC HTB-20) cells were cultured in Richter’s Modification MEM (IMEM) (Corning), supplemented with 10% FBS, 1% P/S, and 20 μg/mL insulin; these cells were maintained at 37° C in a 5% CO_2_ atmosphere(57).

MDA-MB-231 (ATCC HTB-26) cells were cultured in Dulbecco’s Modified Eagle Media (DMEM) (Gibco) supplemented with 5% FBS and 1% P/S; these cells were maintained at 37° C in a 5% CO_2_ atmosphere(58).

To facilitate the use of fluorescent microscopy and automated cell quantification, stable fluorescent cell lines were established (MCF7-EGFPNLS1, BT474-GNS2, 231-GNS) expressing constitutive EGFP with a nuclear localization signal. Genomic integration of the EGFP expression cassette was accomplished utilizing the Sleeping Beauty transposon system. The EGFP-NLS sequence was obtained as a gBlock (IDT) and cloned into the optimized Sleeping Beauty transfer vector pSBbi-Neo. pSBbi-Neo was a gift from Eric Kowarz (Addgene plasmid #60525)(59). To mediate genomic integration, this two-plasmid system consisting of the transfer vector containing the EGFP-NLS sequence and the pCMV(CAT)T7-SB100 plasmid containing the Sleeping Beauty transposase was co-transfected into the MCF-7 population utilizing Lipofectamine 2000. mCMV(CAT)T7-SB100 was a gift from Zsuzsanna Izsvak (Addgene plasmid # 34879)(60). Following gene integration with Sleeping Beauty transposase, eGFP+ cells were collected by fluorescence activated cell sorting and maintained in media supplemented with 200 ng/mL G418 in place of P/S.

### Doxorubicin Treatment

Doxorubicin hydrochloride (Cayman Chemical 15007) is reconstituted in water. Cell culture media is replaced with complete growth media containing doxorubicin at the specified concentration. After 24 hours, doxorubicin media is replaced with drug-free media.

### Long term doxorubicin response experiments

Cells are seeded in a 96-well plate at target densities of 2,000 cells per well. Fluorescent and phase contrast images are collected at intervals of 4 hours or shorter throughout the study in the Incucyte S2 Live Cell Analysis System (Essen/Sartorius) with temperature and environmental control. Cells are initially seeded in 100 μL growth media per well and cultured for 2 days to allow cell adhesion and recovery from the passaging process. Drug treatment is performed by adding 100 μL growth media containing doxorubicin at 2x the desired final concentration, with 6 or 12 replicates for each experimental condition. After 24 hours, the drug exposure is ended by aspirating the media, and replacing with fresh growth media. Images collection is continued for 21-56 days at a frequency between once per two hours and once per four hours; the duration of image collection is selected such that all wells in which any cells recover proliferation are allowed to reach the logistic growth stage.

### Image Analysis

Cell counting is performed using the green fluorescence channel using standard image analysis techniques: background subtraction, followed by thresholding, edge detection, and a minimum area filter. These settings have been optimized for accuracy and robustness in handling the image-to-image variation in acquired images.

### Data normalization

The data for each replicate culture is truncated to ensure that only accurate, meaningful data is used for model calibration. Time course cell number data for each replicate culture is truncated at 30,000 cells or when the counted cell number drops by more than 50% as a result of media handling, or when the cell count becomes unreliable as indicated by repeated discontinuities in the cell number vector. In the case of a single discontinuity where fewer than half of the cells are lost, the data is instead normalized to remove the discontinuity; this normalization is performed by specifying that the cell number for time points prior to the discontinuity will be divided by a constant α, calculated such that the first and second derivative of the cell number are smooth across the discontinuity:

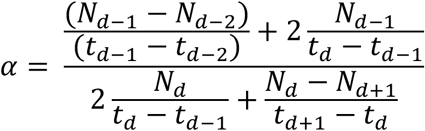

Here *N_d_* is the cell number just after the discontinuity, *N_d−1_* is the cell number just before the discontinuity, and so on.

### Quantification of *t_r_*

To find the proliferation delay, *t_r_*, a density-based clustering technique (DBSCAN) was used to find an initial clustering of cells after treatment had been applied. The optimal parameters for DBSCAN were manually calibrated for each plate. Once a sufficiently large cluster of cells was properly identified, the cluster was tracked backwards in time. At each previous timepoint, DBSCAN was again used to identify the location of the cluster. When a timepoint was identified at which clustering could no longer be found, the proliferation delay was determined as the difference in hours between that timepoint and the time of initial treatment.

### Model calibration

Model parameters are extracted from time course cell population data by calibrating the three models to the cell number vector for each replicate culture using a nonlinear least-squares approach implemented in MATLAB.

### Data Availability

All cell number data in this manuscript, as well as all MATLAB scripts used for model calibration and validation, are available at https://github.com/brocklab/Drug-Response-Dynamics-Model-Calibration.

## Discussion

The use of longitudinal monitoring *via* time-lapse microscopy allowed us to implement and compare dynamic models of drug sensitivity. This allowed us to identify conditions in which a model incorporating a delay on sensitive cell death is preferred over the more common, simpler model, and to determine that across the range of conditions tested, a model describing that delay with an exponential time constant performs best. Additionally, our model selection work identified the parameters *f_r_* and *t_r_* as computationally non-separable. Quantification of *t_r_ via* automated clustering analysis confirmed the existence of a delay in proliferation under some conditions. Incorporating these two time delays removed sources of systematic error and allowed extraction of more accurate values for key model parameters such as the resistant fraction and post-recovery growth rate via calibration.

We identified transient changes in doxorubicin sensitivity following an initial doxorubicin exposure. Inter-treatment interval proved to be an optimizable factor in drug scheduling, with retreatment during the window of transient sensitivity increasing the total time to population doubling by 104% in the MCF7 line, 46% in the BT474 line, and 40% in the MDA-MB-231 line, compared to treatment with an identical total amount of drug with an interval such that drug sensitivity had stabilized. In addition to extending the time to population doubling, treatment at the optimal time actually resulted in elimination of the cancer cell population in 9/12 replicate cultures in the MCF7 cell line and 6/12 replicate cultures in the MDA-MB-231 cell line. The optima identified for these cell lines did not depend on any single model parameter; they result from the interaction of all three relevant parameters, each of which changes dynamically with a different characteristic time.

Although this mathematical modeling-based approach enabled the identification of key features of the dynamic response which have previously not been recognized (i.e., the time delays on both death and proliferation) it does have limitations. While model 1 was selected over the existing bi-exponential model in all three cell lines, the improvement in performance varied with cell line. The calibrated model allowed identification of optimal retreatment intervals in the MCF7 and MDA-MB-231 cell lines, but did not offer a clear equivalent in the BT474 cell line. This is likely due to differences in drug sensitivity and timing of response that characterize each cell line-BT474 has higher sensitivity and responds more slowly; consequently it was treated at a lower dose range. It is likely that the optimal model to describe growth dynamics varies with cell line, and possible that the most relevant mechanisms of changing drug sensitivity vary with cell line. A second significant limitation of this approach is that it is phenomenological, and neither relies on nor contributes to understanding of the underlying mechanisms.

The approach described here allows for high throughput, high time resolution measurement of population-averaged properties. In particular, multidrug treatment would greatly benefit from longitudinal monitoring of population dynamics. Collateral sensitivity, a phenomenon where one drug causes a cell population to become sensitive to a second drug(61), has shown increasing promise *in-vitro*(62,63). Notably, Dhawan et al. found that specific intervals between drugs was necessary in order to identify pairs of collaterally sensitive drugs. Furthermore, the repeatability of collateral sensitivity to drug-resistant populations has yet to be fully understood(64,65). Thus, being able to perform high throughput measurement would allow for pairs of collaterally sensitive drugs to not only be found but evaluated as potential treatments. Additionally, the approach demonstrated here would allow for optimal control theory to be experimentally evaluated, a strategy which can be used to find optimal treatment strategies for a dynamical system(66). Optimal control theory requires a detailed and accurate understanding of the relationship between controllable variables such as drug concentration and inter-treatment interval and biological variables such as the resistant fraction and post-treatment regrowth rate. The experimental and computational methods described here allow for the development of such detailed data sets.

Optimal control theory has recently been used to find optimal scheduling(67,68), but has yet to be evaluated in an *in-vitro* setting. The development of drug response models with sufficient detail for use in predicting collateral sensitivity and testing optimal control theory could additionally contribute to the field of adaptive therapy, in which treatment is modified based on feedback regarding a patient’s individual response(69). This approach currently shows significant promise in the treatment of prostate cancer(70), and the development of more detailed drug response models can contribute to the extension of these techniques to additional cancer types. Single cell sequencing technologies promise to provide great insight into the mechanistic underpinnings of this heterogeneity in drug sensitivity(65,71); however, throughput for these technologies remains low enough (and expense remains high enough) that directly assessing dynamic changes in drug sensitivity over short time scales remains difficult.

These two approaches can complement each other, with longitudinal studies such as the ones described here identifying key time scales and single cell transcriptomic studies enhancing mechanistic insight. This suggests several potentially productive avenues for continuing work: Additional longitudinal studies can extend the techniques demonstrated here to new drugs, new cell lines and new cancer types. Along with the development of new data sets, exploration of additional cancer growth models will allow deeper understanding of what features of a cell line (drug sensitivity, growth rate, clonal heterogeneity, etc.) correspond to particular optimal growth models. The data presented here also indicate time points that would be of particular interest for future single-cell studies, such as the two day post treatment optimal treatment interval in the MCF7 and MDA-MB-231 cell lines. Such studies could elucidate the mechanisms underlying these early changes in drug sensitivity, and could in turn enable another generation of longitudinal studies in cell lines engineered with fluorescent reporters coupled to mechanisms thus identified.

Finally, while these methods are several steps removed from clinically actionable at this point, there is potential to contribute to preclinical drug development. *In vitro* mechanistic studies and preclinical screening often report cell death at a single end point 24-72 hours after treatment; this is insufficient to quantify the long-term drug response of the cell population. Longitudinal studies such as these, implemented iteratively to identify key time scales such as the optimal intertreatment interval, could allow further optimization of drug treatment schedules to inform preclinical testing.

## Conclusions

Using an integrated experimental-computational approach, we identified a biexponential model of drug response including time dependence in post-treatment proliferation and death rates as superior to a static (time independent) model. The delay of proliferation in drug resistant cells is a key variable, and was computationally non-separable from the resistant fraction; consequently, measurement of the proliferation delay is a necessary prerequisite to accurate model calibration. The inter-treatment interval of a multi-dose series was found to be an optimizable parameter of the treatment schedule, increasing the time to a proliferation metric by 40%-106% in three breast cancer cell lines.

## Acknowledgements

We would like to thank Yasmeen K. Zubair and Isha Patel for their work in data collection and processing during the preliminary experiments which initiated this work. We are grateful for funding through the NIH via R01CA226258 and U01CA253540 to A.B. and NCI U01 CA142565, NCI U01 CA174706, NCI U24 CA226110, NCIR01CA240589, and CPRIT RR160005. To T.E.Y. T.E.Y. is a CPRIT Scholar of Cancer Research. The authors declare no conflicts of interest.

